# The central role of the interspecific interactions in the evolution of microbial communities

**DOI:** 10.1101/2022.01.17.476584

**Authors:** Tiffany Raynaud, Manuel Blouin, Marion Devers-Lamrani, Dominique Garmyn, Aymé Spor

## Abstract

The interspecific interactions play an important role in the establishment of a community phenotype. Furthermore, the evolution of a community can not only occur through an evolution of the species composing the community but also of the interactions among them. In this study, we investigated how widespread was the evolution of interspecific interactions in the evolutionary response of eight two-bacterial species communities regarding productivity. We found evidence for an evolution of the interactions in half of the studied communities which gave rise to a mean change of 15% in community productivity as compared to what was expected from the individual responses. Even when the interactions did not evolve themselves, they influenced the evolutionary responses of the bacterial strains within the communities which further affected community response. We found that the evolution within a community often promoted an adaptation of the bacterial strains to the abiotic environment, especially for the dominant strain in a community. Overall, this study suggested that the evolution of the interspecific interactions was frequent and that it could increase community response to evolution. We propose that the existence of an evolution of the interspecific interactions can justify the consideration of the community as a unit of selection.

## INTRODUCTION

What makes a community is the existence of interactions between the species [1, 2]. The interactions give rise to emergent properties at the community-level i.e. properties that are not predictable from the sum of the properties of the component species [3, 4]. It is thus relevant to consider the phenotype of a community as a whole, especially since it is increasingly recognized that community phenotype can respond to the evolution [5]. The evolution of community phenotype can involve or not an evolution of the interspecific interactions. From a theoretical standpoint, it is accepted that microbial community evolution can occur through genetic changes in the community members (e.g. through mutations, horizontal gene transfer, gene loss [6, 7]). Depending on the authors, changes in the relative abundances of the species within a community are also considered as part of the evolution of the community [8, 9] or not [6, 7]. In the latter case they are referred to as “ecological sorting” [10]. Furthermore, as well as the interactions are involved in the establishment of community phenotype, there is also evidence that they can contribute to the evolution of this phenotype. This has been investigated in the field of artificial selection at the community level through modelling [11] and experimental approaches on communities made of two beetle species [12]. Both approaches highlighted that genetic changes in the species within a community are not always sufficient to explain the observed response of the community to selection and suggested that the interspecific interactions can be involved in community evolution. This is likely the strongest argument to support the idea that communities are units of selection.

In parallel, other studies aiming at understanding the evolutionary dynamics of microbial communities, especially in response to environmental changes, provided detailed assessments of the evolution of interspecific interactions in synthetic communities. It has been shown that, in a two-species bacterial community, a mutation in one of the two strains induced a shift from a commensal interaction to a more exploitative one [13]. This shift in the interaction occurred after five days of experimental evolution and gave rise to an enhanced community productivity. It indicates that the interactions can evolve through the evolution of one of the community members. Another way for the interactions to evolve is through the evolution of multiple species in a community. An experimental study showed that, in a four-species bacterial community, changes in resource use in the four species when experimentally evolved in polyculture reduced the occurrence of negative interspecific interactions [14]. It was associated with a higher productivity at the community level than this of a community which was built from the four species evolved in isolation. Changes in the interactions through the evolution of several species can also result from co-evolution, i.e. reciprocal adaptive changes in two populations or species [15]. Coevolution is different from evolution in response to the presence of a species as it involves adaptation and counter-adaptation in the interacting species. In experimental conditions, coevolution can be evidenced either by tracking the evolutionary changes in each of the co-cultured species or by comparing the evolutionary responses in treatments where coevolution is allowed or not [16].

An additional level of complexity emerges from the fact that the evolution of the interspecific interactions can depend on the abiotic environment. For example, in bacterial communities, the evolution of the interspecific interactions can be promoted by a structured environment, allowing the formation of biofilm, as compared to a homogeneous environment [13]. It has also been shown that the involvement or not of the interactions in a bacterial community evolutionary response can depend on the resources or on the pH of the culture medium [10]. Interestingly, it has been shown that the productivity of a community increased as compared to the ancestral community only when the interactions were involved in the community response to evolution (and in the most diverse communities only) [10]. To go further, the abiotic environment can be modified by a species which can influence the evolution of other species in a community, this is referred to as niche construction [17]. It has been shown that the pairwise interaction between a bacteria and a yeast shifted from commensalism to amensalism and then to antagonism when the environment started to be changed by the yeast. Indeed, the excretion of a bacterial growth inhibitor promoted the evolution of resistance in the bacterial population which lowered the fitness of the yeast [18]. Thus, eco-evolutionary feedbacks are also involved in the evolution of the interactions and of the communities.

There are many studies that illustrate well the evolution of the interspecific interactions, the question is not anymore *whether the interactions can evolve* but *how important is the evolution of the interactions in the communities* [7]. In this study, we aimed at providing an insight into the prevalence of the evolution of interactions in the evolution of community phenotype. Following a five-month experimental evolution of synthetic bacterial communities [19], we re-isolated eight pairs of strains that evolved in different communities. We assessed community and community member phenotypes by measuring the optical density as a proxy of productivity. We compared them with the ancestral phenotypes and the phenotypes obtained by assembling the same strains evolved in isolation to discuss the evolution of interactions. We hypothesized that: *i)* the interspecific interactions played a role in the evolution of community phenotype (i.e. the phenotype of the evolved community would be different from this of a community reconstructed from strains that evolved in isolation); *ii)* this role occurred through an evolution of the interactions themselves (i.e. the evolutionary response of the community is not predictable from the evolutionary responses of the community members); *iii)* the evolution of the community phenotype depended on the abiotic environment. To verify this third hypothesis, we assessed the phenotype of the evolved strains and communities in a second abiotic environment in order to discuss the adaptation to the abiotic conditions of the environment of the experimental evolution.

## MATERIALS AND METHODS

### Origin of the studied communities

The eight two-strain communities studied in this experiment stemmed from an experimental evolution procedure in which bacterial strains (= monocultures) and communities were grown for five months with a serial transfer each 3.5 days. This experiment involved 18 laboratory strains that were used to create communities differing for their initial richness levels (see [19]). During the experimental evolution, the strains and communities were grown in sterile 2 ml deep-well plates (Porvair Sciences, Wrexham, UK) filled with 1 ml of a mix of 1:5 lysogeny broth (LB) and 1:5 tryptic soy broth (TSB), hereafter called EE medium for Experimental Evolution, and placed at 28°C without shaking. An optical density (OD) measurement was performed at each serial transfer (as a proxy of productivity) and the transfer occurred following two treatments: artificial selection (where the transferred replicate was the one with the highest OD among ten) and no artificial selection (where the replicate was transferred whatever its OD). The monocultures and communities were stored at −80°C in 30% glycerol before the experimental evolution (ancestors) and after the experimental evolution (evolved strains and communities). In a first step of isolation, all of the communities of richness level 2 (six), both under artificial selection and no artificial selection (see [19]), were considered for being analysed in the present study. In a second step, all of the communities of richness level 4 (six) either under artificial selection or under no artificial selection were also considered to complete the experimental design. The pairs of strains that were finally included in the experiment are presented in Table 1 and responded to the following criteria: successful isolation of the strains from the evolved community and availability of the corresponding strains evolved in isolation.

**Table 1.** Two-strain communities studied in the experiment. Some of the pairs of strains evolved in the absence of other strains (i.e. in two-strain native communities), whereas other pairs evolved in the presence of other strains (i.e. in four-strain native communities), this is specified into the column “Initial richness level of the native community”. Some of the native communities evolved under artificial selection whereas others evolved under “no artificial selection” (i.e. natural selection only), this is specified in the column “Selection regime applied to the native community”. In each community, strain 1 is the most productive of the two strains (highest OD) and strain 2 is the least productive one (lowest OD).

| Community identifier | Strains | Initial richness level of the native community | Selection regime applied to the native community |
| --- | --- | --- | --- |
| Community A | 1 *Variovorax* sp.38R<br>2 *Pseudopedobacter saltens* DSM1 2145 | 2 strains | No artificial selection |
| Community B | 1 *Variovorax* sp.38R<br>2 *Pseudopedobacter saltens* DSM1 2145 | 4 strains | No artificial selection |
| Community C | 1 *Pseudomonas knackmussii* DSM6978<br>2 *Variovorax* sp.38R | 4 strains | No artificial selection |
| Community D | 1 *Pseudomonas* sp. ADPe<br>2 *Escherichia coli* WA803 | 2 strains | No artificial selection |
| Community E | 1 *Pseudomonas knackmussii* DSM6978<br>2 *Pseudopedobacter saltens* DSM1 2145 | 4 strains | No artificial selection |
| Community F | 1 *Pseudomonas* sp. ADP3<br>2 *Escherichia coli* K12 | 4 strains | No artificial selection |
| Community G | 1 *Escherichia coli* WA803<br>2 *Agrobacterium* sp.9023 | 4 strains | Artificial selection |
| Community H | 1 *Pseudomonas* sp. ADPe<br>2 *Escherichia coli* WA803 | 2 strains | Artificial selection |

### Isolation of the strains from the evolved communities

To isolate the strains that evolved in communities, we revived the evolved communities from the glycerol stocks by growing them on agar plates (EE medium) by streaking. After 72h of growth at 28°C, we picked the colonies of differing morphologies and placed them on new separated agar plates by streaking. After a new cycle of growth, one colony per plate was picked, placed in 200 μl of 0.9% NaCl and 100 μl of this suspension were plated on an agar plate with glass beads. At this step, 2 μl of suspension were used to perform a PCR for the identification of the strains (see below). After a new cycle of growth, several colonies were picked on each plate and put in 20 ml of medium in a flask (48h, 120 rpm). 800 μl of suspension were then stored at −80°C in 800 μl of 60% glycerol. As these isolation steps required four growth cycles during which evolution could act, we also performed these four growth cycles in the same conditions for the corresponding ancestral and evolved in isolation strains.

### Identification of the strains

A PCR targeting 16S rRNA gene with the primers 27F/1492R [20] was performed for each strain isolated from the evolved communities. A digestion of the PCR products was then performed with the *AluI* restriction enzyme and followed by an electrophoresis for the identification of the strains at the genus level. For the genera that were represented by several strains in our experiment (i.e. *Pseudomonas* and *Escherichia*), we performed further analyses for an identification at the strain level. We used data from *gyrB* sequencing at the community level to determine which *Pseudomonas* strain was present in the community and coupled it with analyses at the strain level for formal identification. The different strains were identified based on the presence or not of *atzD* gene (assessed by PCR) and the resistance or not to nalidixic acid and amoxicillin (assessed by growing the strains on agar plates containing a mix of the two antibiotics at a final concentration of 100 μg.ml^-1^). *Escherichia coli* K12 and *Escherichia coli* WA803 were identified based on their ability to do or not lactose fermentation (which was assessed by growing the strains on agar plates on Drigalski agar medium).

### Evolutionary history treatments

Each of the two strains of a community (eight in total, hereafter identified as communities A to H; Table 1) was grown in its ancestral version (i.e. before experimental evolution), in its “evolved in isolation” version (i.e. after experimental evolution as an isolated strain) and in its “evolved in community” version. It resulted in six treatments (two strains and three evolutionary histories per strain; Figure 1a). Within each community, the most productive (highest OD at 3.5 days) of the two ancestral strains was referred to as “strain 1” and the least productive was referred to as “strain 2”. In addition, each community was grown in its ancestral version (i.e. co-culture of the two ancestral strains), in its “evolved in isolation” version (i.e. co-culture of the two strains that evolved in isolation) and in its “evolved in community” version (i.e. co-culture of the two strains that evolved together in a community). Two communities mixing ancestral strains and strains evolved in community were also included: mixed community 1 (i.e. co-culture of strain 1 evolved in community and ancestral strain 2) and mixed community 2 (i.e. co-culture of strain 2 evolved in community and ancestral strain 1). It resulted in five treatments at the community level plus the six treatments at the strain level (Figure 1b); each treatment was replicated eight times per each studied community.

**Figure 1.**
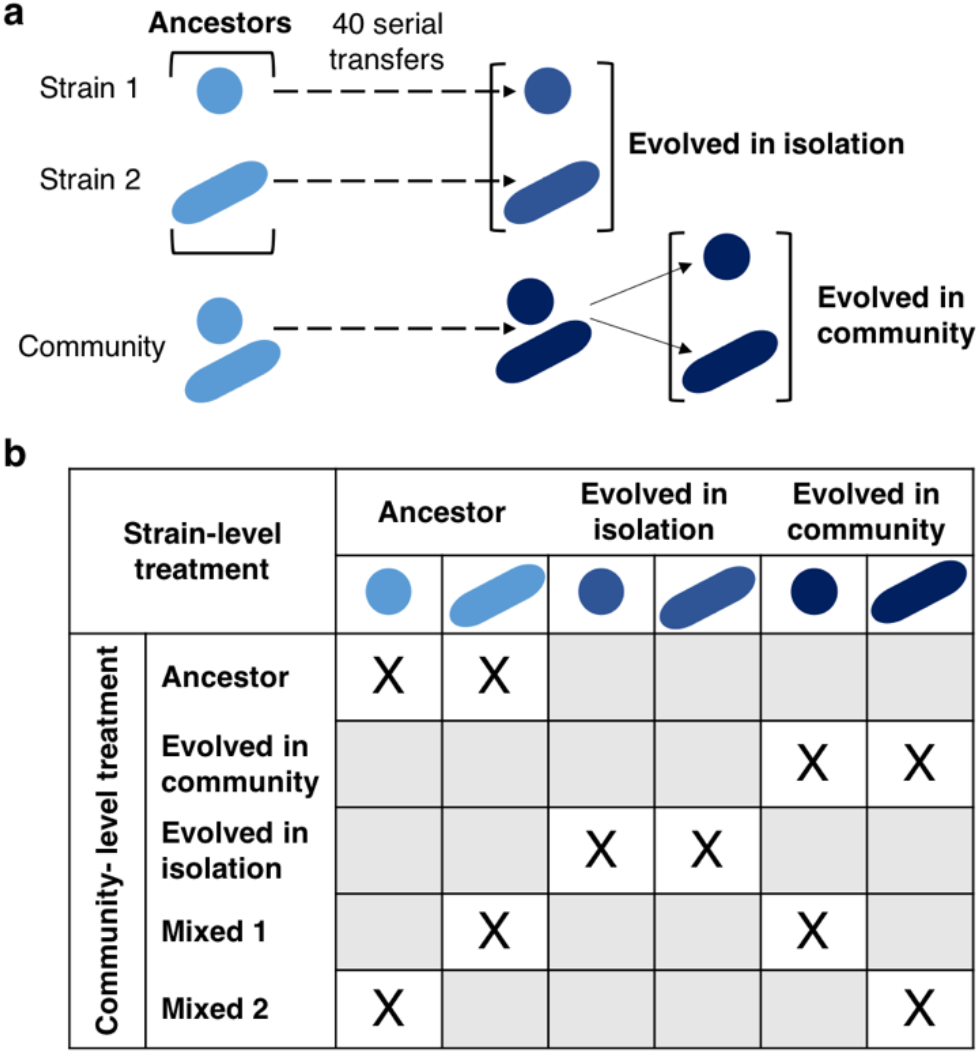
Experimental design. **a)** Each bacterial strain was previously experimentally evolved in isolation and as a member of a community [19]. At the end of this experimental evolution, the strains were isolated from the community in which they evolved. **b)** From the strains, different communities were built: ancestor (co-culture of two ancestral strains), evolved in community (co-culture of two strains that evolved together), evolved in isolation (co-culture of two strains that evolved in isolation), mixed 1 (co-culture of one ancestral strain and one strain evolved in community) and mixed 2 (co-culture of one ancestral strain and one strain evolved in community conversely to mixed 1).

### Community construction, growth conditions and phenotype assessment

Before the start of the experiment, each strain was revived from the glycerol stock and grown in 20 ml of EE medium in a flask (48h, 28°C, 110 rpm). The OD of the suspensions was measured (Infinite M200 PRO, Tecan, Männedorf, Switzerland) and the suspensions were diluted to a final OD of 0.002 in EE. The eight two-strain communities were built by mixing an equivalent volume of each of the suspensions of the required strains. Then, two plates per community were inoculated with the suspensions at OD 0.002: a 2 ml deep-well plate (1 ml of suspension per well, eight replicates per treatment) and a honeycomb plate (Thermo Fisher Scientific, Waltham, Massachusetts, USA; 400 μl of suspension per well, eight replicates per treatment). The growth conditions in deep-well plates were: 28°C, no shaking; the OD was measured after 3.5 days of growth by homogenising the well content, pipetting 200 μl of suspension and transferring it into a new plate for OD measurement at 600 nm (Infinite M200 PRO). These growth conditions were identical to the growth conditions of the experimental evolution, hereafter we refer to these conditions as “environment 1”. The growth conditions in honeycomb plates were: 28°C, 15 s of shaking 5 s before each OD measurement (600 nm; Bioscreen, Oy Growth Curves Ab Ltd, Helsinki, Finland), one measurement every 30 min. These growth conditions were different from these of the experimental evolution, hereafter we refer to these conditions as “environment 2”.

### Statistical analyses

The OD after 3.5 days of growth was analysed in two steps with two linear mixed models. The following model was used to analyse the effect of the evolution on strain and community phenotypes:

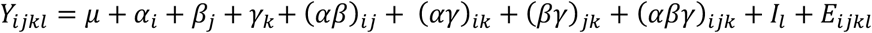

*Y_ijkl_* is the OD of the biological entity *i* (three levels: strain 1, strain 2, community), of identity *l* (24 levels: strain or community identity), of evolutionary history *j* (three levels: ancestor, evolved in isolation, evolved in community), in environment *k* (two levels: environment 1, environment 2). *μ* is the intercept, *α_i_* is the effect of the biological entity, *β_j_* is the effect of the evolutionary history, *γ_k_* is the effect of the environment. The interaction effects between *i)* the biological entity and the evolutionary history (*αβ*)*_ij_*; *ii)* the biological entity and the environment (*αγ*)_*ik*_; *iii)* the evolutionary history and the environment (*βγ*)*_jk_*; *iv*) the biological entity, the evolutionary history and the environment (*αβγ*)*_ijk_* were also included in the model. *I_l_* is the random effect of the strain or community identity, *E_ijkl_* is the residual error.

A second linear mixed model was built to analyse the effect of the evolutionary history of the community members on the community phenotype:

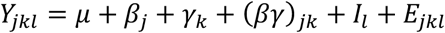

*Y_jkl_* is the OD of the community of identity *l* (8 levels: A to H), of evolutionary history *j* (five levels: ancestor, evolved in isolation, evolved in community, mixed 1, mixed 2), in environment *k* (two levels: environment 1, environment 2). *μ* is the intercept, *β_j_* is the effect of the evolutionary history, *γ_k_* is the effect of the environment, (*βγ*)*_jk_* is the effect of the interaction between the evolutionary history and the environment. *I_l_* is the random effect of the community identity, *E_jkl_* is the residual error.

To go into the details of the responses for each community, the OD after 3.5 days was then analysed with a linear model that included the identity of the individual as a fixed effect factor as well as the evolutionary history and the interaction between the identity and the evolutionary history. One model was built for the strains and one for the communities in both environments. Then, the predictability of the community evolutionary response was analysed by comparing the response of the community (i.e. change in OD during experimental evolution) to *i)* the response of strain 1 evolved in community, *ii)* the response of strain 2 evolved in community *iii)* the sum of the responses of strains 1 and 2 (which corresponds to the expected response under the hypothesis of an additivity of the individual responses, i.e. the absence of evolution of interspecific interactions). The mean responses and the corresponding 95% confidence intervals were obtained by bootstrapping (1 000 iterations of the calculation of the response from randomly sampled values with replacement).

All the analyses were performed with R software version 3.6.3 with lmerTest package for linear mixed models [21], car package for type II analyses of variance [22] and emmeans package for pairwise comparisons [23].

## RESULTS

### The strains’ responses are driven by their initial productivity in monoculture

The effect of the evolutionary history on optical density (OD) depended on the biological entity, i.e. whether the considered phenotype was this of the community or of the community members, and it also depended on the environment (biological entity*history*environment: *χ*^2^=48; p_df=4_=1.0×10^−9^; Table S1). Strain 1 and strain 2, the initially most and least productive strain respectively, responded differently to the evolution in environment 1. The OD of strain 1 when evolved in community tended to be higher than this of strain 1 as an ancestor and was higher than strain 1 evolved in isolation (respectively 0.68±0.18, 0.66±0.12, 0.55±0.22; Figure 2). On the contrary, strain 2 showed a lower OD when evolved in community as compared to evolved in isolation (0.30±0.12 and 0.37±0.18 respectively).

**Figure 2.**
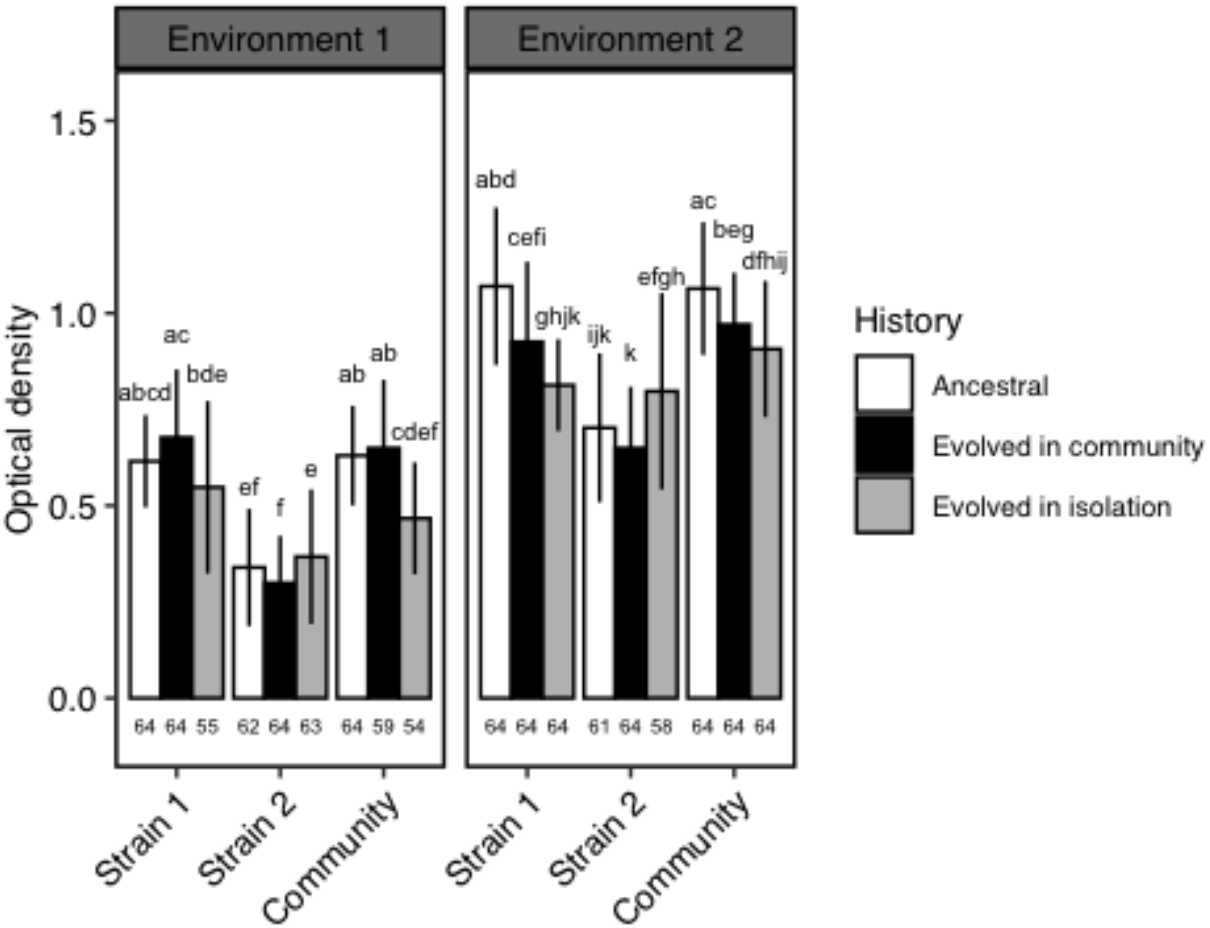
Optical density of the community and the community members depending on the evolutionary history and the environment. Environment 1: identical growth conditions to the experimental evolution; Environment 2: different growth conditions from the experimental evolution. Strain 1 is the most productive of the two strains in a given community (highest OD) and strain 2 is the least productive of the two strains (lowest OD). Community refers to the co-culture of strain 1 and strain 2. Different letters represent significant differences in OD within a given environment. Mean values are given ± SD. Sample sizes are given on the bottom of the graphs.

### Community response is driven by the most productive strain in monoculture

In environment 1, the OD of the communities composed of strains that evolved together was not significantly different from the OD of the ancestral communities (respectively 0.65±0.18 and 0.63 ±0.13; Figure 2). But, it was higher than the OD of the communities in which the members evolved in isolation (0.47±0.15) suggesting that the evolution in community (i.e. co-culture) did not produce the same outcome than an evolution in isolation. However, the communities composed of strains that evolved together produced the same phenotype as mixed communities (one ancestral and one evolved strain, Figure 3).

**Figure 3.**
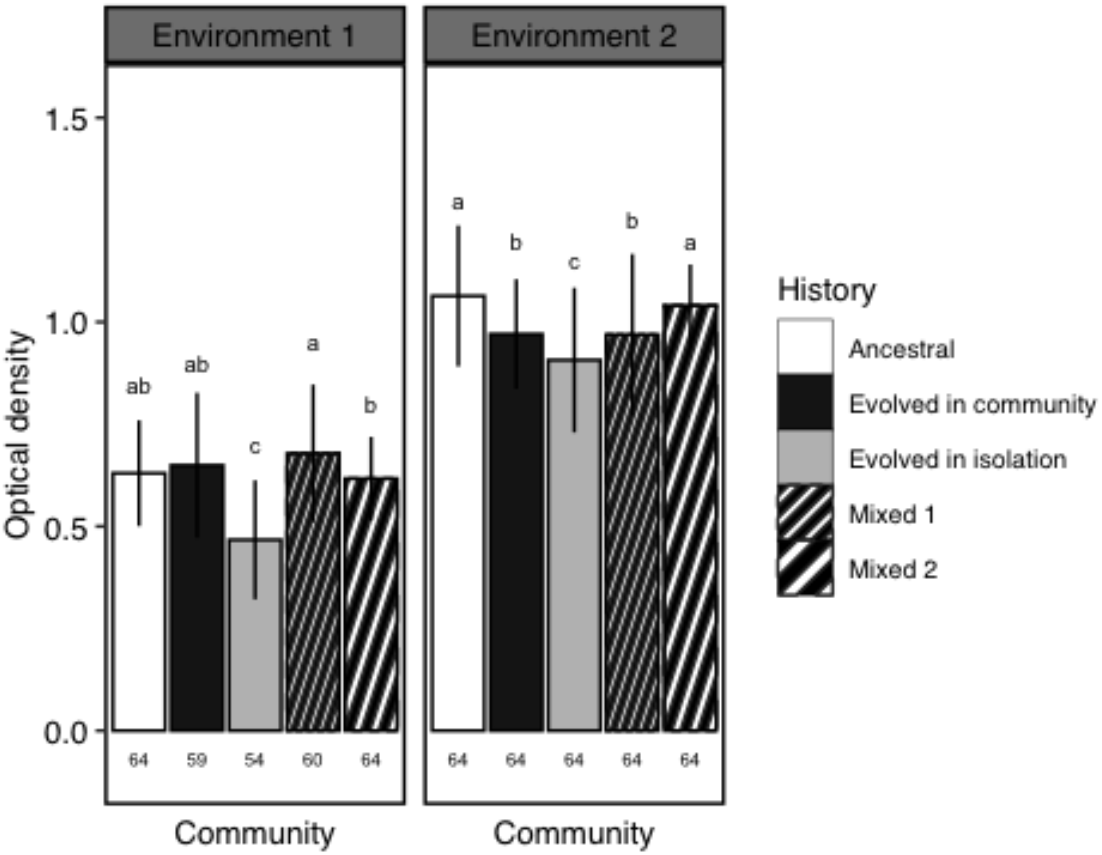
Optical density of the communities depending on the evolutionary history of their members and the environment. Environment 1: identical growth conditions to the experimental evolution; Environment 2: different growth conditions from the experimental evolution. Community refers to the co-culture of strain 1 and strain 2. In mixed community 1, the strain 1 evolved in community was grown with the ancestral strain 2. In mixed community 2, the ancestral strain 1 was grown with the strain 2 evolved in community. Different letters represent significant differences in OD within a given environment. Mean values are given ± SD. Sample sizes are given on the bottom of the graphs.

The OD of the community was not different from the OD of strain 1 whatever the evolutionary history (respectively 0.61±0.16 and 0.62±0.18 on average; Figure 2). Also, the response of the community to the evolutionary history was similar to this of strain 1 (i.e. trend to increase in OD with an evolution in community as compared to the ancestor and trend to decrease in OD with an evolution in isolation; Figure 2). Thus, community phenotype seemed to be driven by strain 1.

### Community response involves an evolution of the interactions in half of the cases

For all of the studied communities, there was a significant difference in OD between the evolved in community and evolved in isolation treatments (Figure 4) which highlighted the importance of the interactions in the evolution of community phenotype. This difference was in favour of the evolved in community treatment in seven of the eight communities (Figure 4). One evolved community showed no difference in OD as compared to the ancestral community (community G; Figure 4). Three communities showed differences in OD with the communities of all other evolutionary histories (communities A, C and F). It indicated that, in these cases, the only way to obtain the evolved community phenotype was through the presence of the two strains in their evolved in community version. The four remaining communities (B, D, E and H) showed no difference in OD as compared to at least the mixed community 1 (Figure 4) highlighting the role of strain 1 in the expression of the evolved community phenotype in these cases.

**Figure 4.**
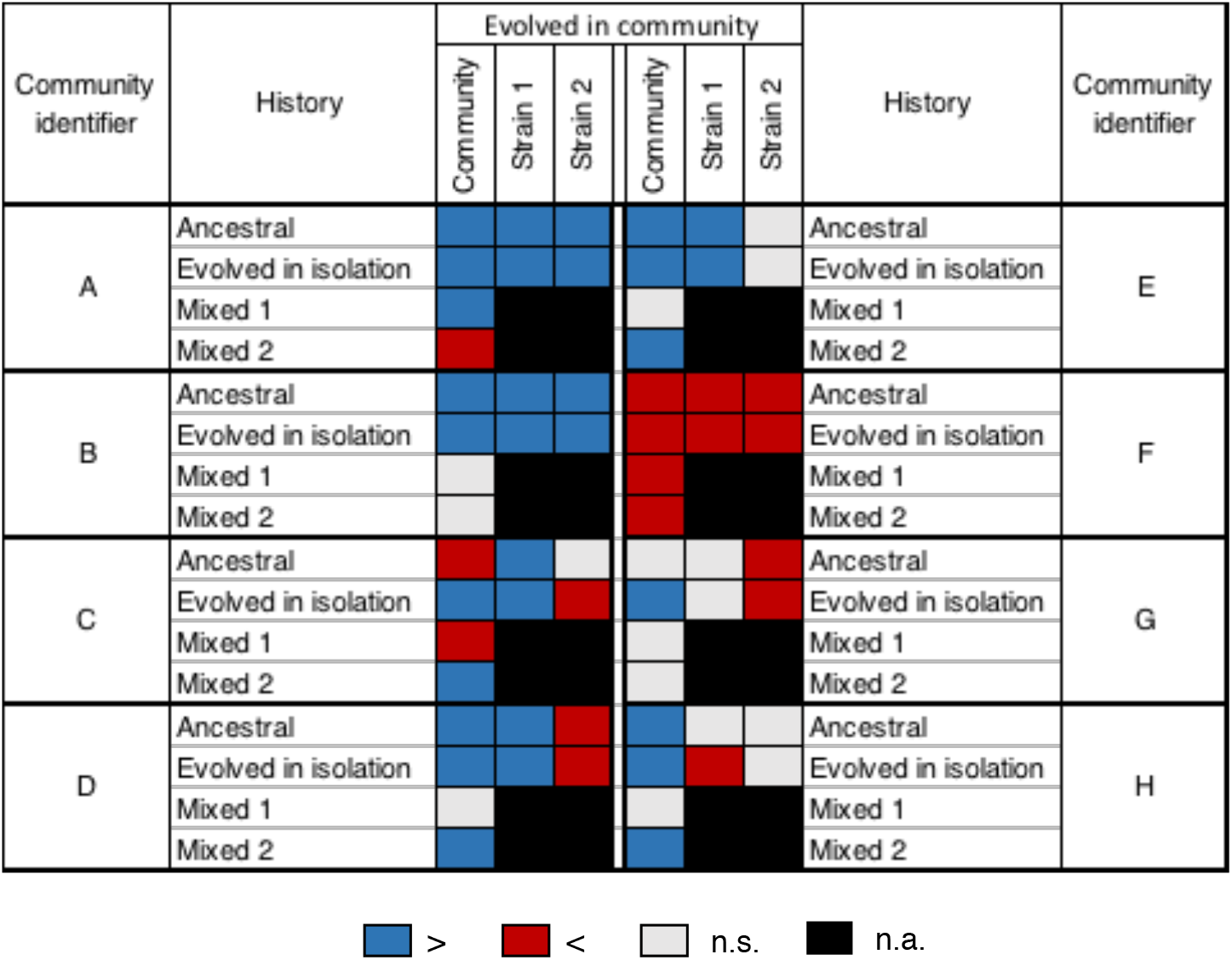
Effect of an evolution in community on optical density in environment 1 depending on the community. The OD of a community composed of strains that evolved together (in columns) is compared to the OD of a community including ancestral strains, strains evolved in isolation or one ancestral strain and one strain evolved in community (mixed 1 and mixed 2) (in rows). The OD of strains 1 and 2 evolved in community (in columns) is compared to the OD of the corresponding strain as an ancestor or evolved in isolation (in rows). Blue: significantly higher. Red: significantly lower. Light grey: no significant difference (α=0.05). Black: not applicable.

The evolutionary response of four of the communities was predictable neither from the responses of the community members nor from the expected response under the hypothesis of an additivity of the individual responses, i.e. an absence of evolution of the interspecific interactions (communities A, C, F and H; Figure 5). It suggested that the evolutionary response involved an evolution of the interactions. In communities D and E, the community response was predictable from the response of strain 1 and in community B, it was predictable from the sum of the responses of the two strains (Figure 5). Thus, we did not evidence an evolution of the interspecific interactions in these communities.

**Figure 5.**
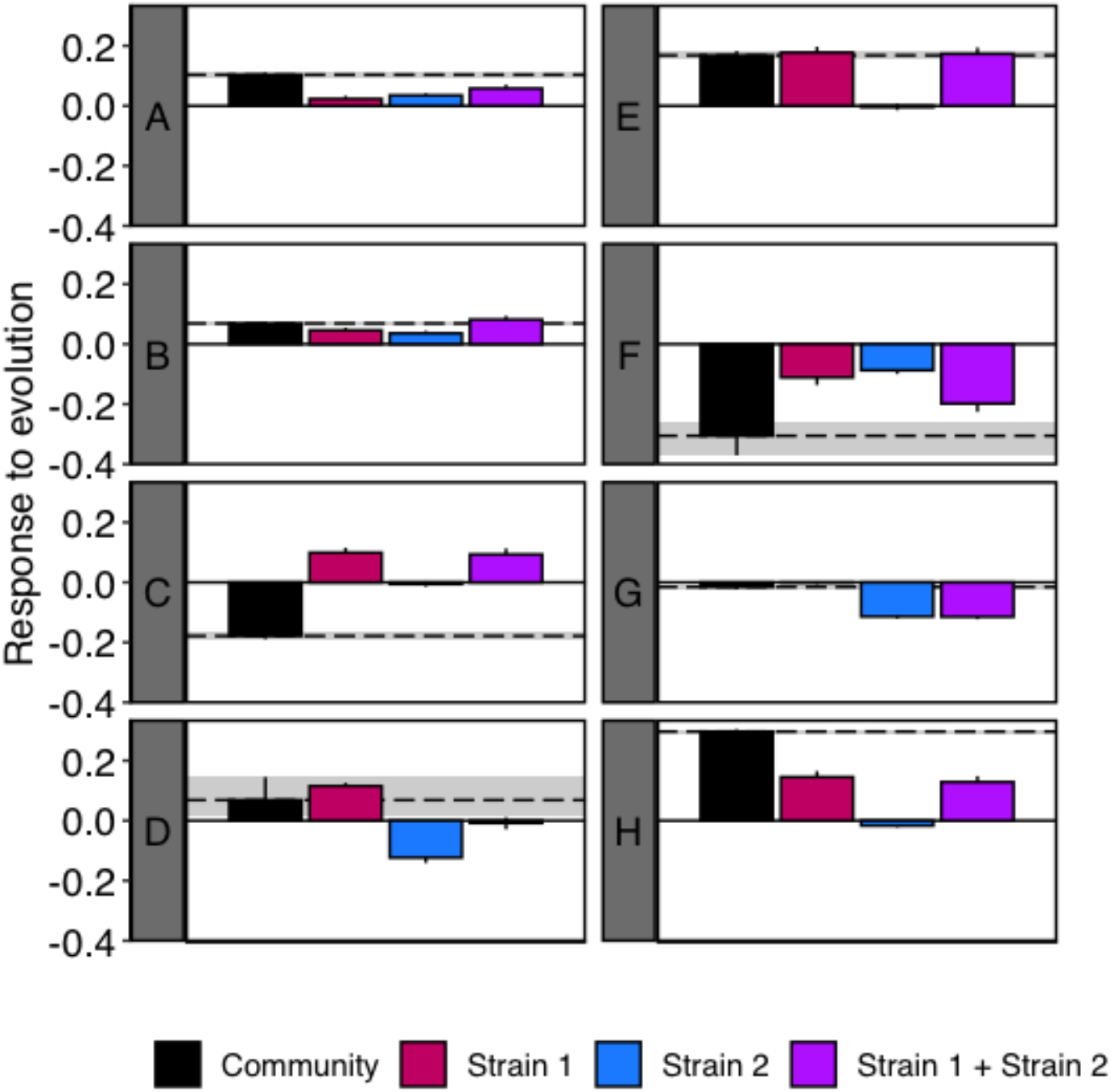
Predictability of the evolutionary response of the community. The observed evolutionary responses of the community, strain 1 and strain 2 in environment 1 were expressed as the difference in optical density as compared to the corresponding ancestor (i.e. ancestral community or ancestral strain 1 or ancestral strain 2 in environment 1). “Strain 1 + Strain 2” refers to the expected response to evolution under the hypothesis of an additivity of the individual responses, it was obtained by summing the observed responses of strain 1 and strain 2. Bars represent 95% CI. On each graph, the black dashed line represents the mean value of the response of the community and the grey area represents the associated 95% CI. Means and 95% CI were obtained by bootstrapping.

### The abiotic environment influences the evolutionary responses

In environment 2, where the conditions differed from these of the experimental evolution, strain 2 showed a similar response to the evolutionary history than in environment 1 (Figure 2). On the contrary, the responses of strain 1 and of the community changed: the highest OD was observed for the ancestors followed by evolved in community and by evolved in isolation treatments. The expression of the “evolved phenotype” thus depended on the abiotic environment. As in environment 1, community phenotype and community response to the evolutionary history were similar to strain 1 (Figure 2). And, the OD of the mixed community 1 was similar to this of the community in which the strains evolved together (respectively 0.97±0.20 and 0.97±0.13) whereas mixed community 2 showed a higher OD that did not differ from this of the ancestral community (Figure 3), again highlighting the influence of strain 1 on community phenotype.

Going into the details of the responses of the different communities, some of the changes in the strain and community phenotypes were consistently observed whatever the environment whereas others were not detectable or occurred in the opposite direction when the environment changed (Figure 6a). The phenotypic change in response to evolution in the evolved community (i.e. change in OD as compared to the ancestral community or to the community with evolved in isolation members) was maintained in environment 2 for three communities over eight (A, C and F; Figure 6b). When a strain that evolved in community showed a significant increase in OD as compared to the ancestor, this pattern was always lost when the environment changed (Figure 6c and d). On the contrary, when a strain that evolved in community showed a significant decrease in OD as compared to the ancestor, this pattern was maintained in environment 2 in three cases over four. The changes in OD in the strain that evolved in community as compared to the corresponding strain evolved in isolation were maintained in environment 2 in nine cases over 13 (Figure 6c and d).

**Figure 6.**
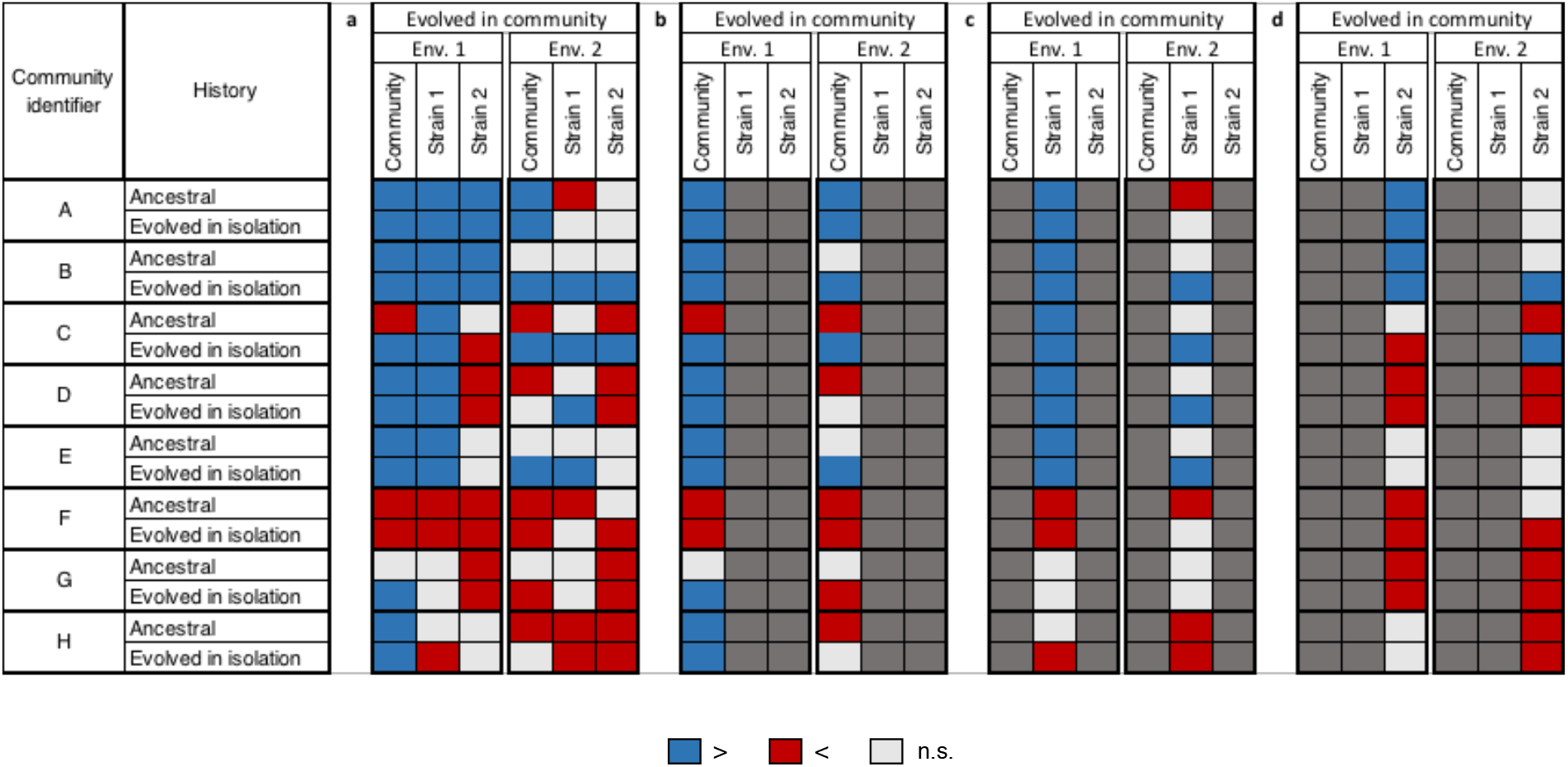
Effect of the environment on the expression of the evolved phenotype. The OD of a community composed of strains that evolved together (in columns) was compared to the OD of a community including ancestral strains or strains evolved in isolation (in rows). The OD of strains 1 and 2 evolved in community (in columns) is compared to the OD of the corresponding strains as ancestors or evolved in isolation (in rows). The results are presented for both environments. Environment 1: identical growth conditions to the experimental evolution; Environment 2: different growth conditions from the experimental evolution. Blue: significantly higher. Red: significantly lower. Light grey: no significant difference (α=0.05). The overall results are shown on panel a, and panels b, c, and d show the results of the community, strain 1 and strain 2 respectively. On those panels, only the comparisons of interest are shown, and the others are shaded in dark grey for readability.

## DISCUSSION

We showed that the evolution of the strains in a community was influenced by the interspecific interactions. Indeed, an evolution in isolation did not produce the same phenotype as an evolution in community (Figure 2). These results are in accordance with an increasing body of literature that highlights the effect of the biotic context, i.e. of the evolution within a community, on the evolutionary response and on the fitness of the community members [10, 13, 24, 25]. The characterization of the community members on the basis of their productivity before experimental evolution allowed a good explanation of their responses to evolution, despite the fact that we grouped species from different genera under the entities “strain 1” and “strain 2” (model R^2^=0.85; Table S1). To go further, we found that the most productive strain had a dominant role in explaining community phenotype and community response to evolution (Figure 2). It was probably highly linked to the fact that the studied community phenotype was productivity but, it also suggested that the most productive strain in monoculture was also the dominant strain in the community as previously observed on two-species communities [26].

Beyond an effect at the individual level, our results indicated that the evolution of community phenotype, i.e. productivity, was influenced by evolutionary changes in interspecific interactions. Indeed, as in a previous study [14], the phenotype of the evolved community could not be obtained by reconstructing a community from strains that evolved in isolation (Figures 2 and 4). We observed an effect of the interactions on community evolutionary response in all of the communities that showed an evolution in their phenotypes, i.e. seven among the eight (Figure 4, except G). However, this effect of the interactions depended on the studied community and occurred through three different ways. Community phenotype evolved through *i)* an evolutionary response of one strain conditionally to the presence of the second strain without evolution of the interaction (communities D and E), *ii)* an evolutionary response of the two strains conditionally to their respective presence without evolution of the interaction (B), *iii)* an evolution of the interaction itself under the influence of one (H) or of the two strains (A, C and F; Figures 4 and 5). Thus, the evolution of the community phenotype involved an evolution of the interactions in more than half of the cases. It suggested that the implication of the evolution of the interactions in the evolution of community phenotype is not rare in experimental evolution of microbial communities. In another study [11], a modelling approach allowed to estimate that the responses of ecosystems to evolution under artificial selection would involve an evolution of the interspecific interactions in 4% of cases when targeting an increase in a property and in 38% of the cases when targeting a decrease in a property (this could be modulated by specific experimental choices). More recently, it has been estimated that the evolution of the productivity of beech tree bacterial communities was explained by ecological sorting at 0.35%, by additive evolution at 17.7% and by the evolution of the interspecific interactions at 14.3% [10]. It is not straightforward to estimate the importance of the interspecific interactions in community evolutionary dynamics as their role seems to be highly dependent on the studied community but, together, these results suggest that it is relevant to consider the evolution of the interactions when studying community dynamics, at least in laboratory experiments. Since interspecific interactions, which are not definable below the level of the community, are potential determinants of the evolution of a community, it can be necessary to consider the community as a selection unit.

In the communities in which an evolution of the interspecific interactions was detected, the change in community productivity was higher than expected but the direction of this change was community-dependent. The response to evolution when the interactions evolved (i.e. in communities A, C, F and H) gave rise to a mean change in productivity of 35±13%, i.e. +15±7% as compared to what was expected from the individual responses. However, in two communities over four (C and F) this change was negative (i.e. the productivity of the evolved community was lower than this of the ancestral community) and in one case it occurred whereas the sum of the individual responses was positive (C; Figure 5). In the other studies that reported an evolution of the interactions, the effect was to enhance community productivity [10, 13, 14]. Furthermore, some authors registered a reduction of the negative interactions and the evolution towards positive ones [14]. In our study, we did not characterize the interactions, but we can hypothesize that different types of interactions (i.e. positive or negative) led to different responses of the community phenotype to the evolution of the interactions.

The influence of the abiotic environment on the evolutionary responses of the communities and community members was community-dependent. For three of the four communities in which an evolution of the interactions was detected, the response to evolution was consistently observed in the two environments (communities A, C and F; Figure 6b) contrary to what was observed for the strains composing these communities (Figure 6c and d). A possible explanation would be that the evolutionary responses of the strains involved an adaptation to the abiotic component (so that the response is not consistently observed when changing the environment) but that the expression of the “evolved” interaction did not rely on an adaptation to the abiotic component or relied on an adaptation to a condition that is found in the two environments [27]. Previous studies have shown the importance of the resources on the outcome of the evolution of interactions [14, 28]. As the same culture medium was used in the two environments in our experiment, it could suggest that the evolution of the interactions implied modifications in resource sharing.

Our results also suggested that the evolution in community often promoted an adaptation of the strains to the abiotic component, especially in strain 1 (Figure 6c and 6d). This is not expected since the theory predicts that there are trade-offs between the adaptation to the abiotic and to the biotic components [6, 14] and, that biotic forces are dominant over abiotic forces in driving species evolution (Red Queen hypothesis; [29]). Thus, it is expected that strains that evolved in isolation would show a better adaptation to the abiotic environment than strains that evolved in community. It has been observed experimentally [14, 30] but seemed to be strain-dependent. Our results suggested that the interspecific interactions could have promoted evolutionary responses to the abiotic conditions, which can occur through competition for example [6]. These results may be linked to the structure of the environment. Indeed, it has been suggested that in homogeneous environments, the evolution would act through the selection of traits that are directly beneficial for the carrier species [7]. Thus, the evolutionary response of a strain to the presence of another strain could be an adaptation to the abiotic conditions, which could have a direct and positive effect on the strain fitness.

In this study, we aimed at investigating the importance of the evolution of the interactions in community evolution. There was evidence for an evolution of the interactions in half of the studied communities. Moreover, we highlighted that, even when they did not evolve themselves, the interactions influenced the evolution of both community phenotype and community members’ phenotype. It is thus relevant to consider that a community is not just an addition of evolving species, but that the community by itself, with evolving species interactions, can be the unit of selection. The present study included eight communities, eight bacterial strains belonging to five genera and focused on pairwise interactions only. Further studies involving higher levels of community complexity are needed to investigate how widespread is the importance of interspecific interactions in community evolutionary dynamics. To go further, our results suggested that the communities in which the interspecific interactions evolved were more likely to be independent on the abiotic environment to express the evolved community phenotype. This is of particular interest in the field of the artificial selection at the community level and its possible applications.

## Acknowledgements

We thank Alain Hartmann and Baptiste Serbource, MERS team, INRAE Dijon, for their advice and the provision of the equipment allowing the assessment of the antibiotic resistance of the strains.

