## Supplemental Table 1 for "The central role of the interspecific interactions in the evolution of microbial communities"

**Table S1 Analysis of variance (ANOVA) of the optical density (OD) of the communities and community members.** The effect of the biological entity (strain 1, strain 2, community), the history (ancestors, evolved in community, evolved in isolation), the environment (1, 2) and their interactions on OD were estimated with a linear mixed model including the identity of the strain or community as a random effect factor. The conditional $R^2$ is presented (i.e. variance explained by both fixed and random effect factors; the marginal $R^2$ – fixed effect factors only – was 0.63).

| | Df | Chi squared | p |
|---|---|---|---|
| Biological entity | 2 | 19.3 | **$6.38\text{x}10^{-5}$** |
| History | 2 | 104 | **$<2.2\text{x}10^{-16}$** |
| Environment | 1 | 2817 | **$<2.2\text{x}10^{-16}$** |
| Biological entity * History | 4 | 193 | **$<2.2\text{x}10^{-16}$** |
| Biological entity * Environment | 2 | 19.3 | **$6.32\text{x}10^{-5}$** |
| History * Environment | 2 | 46.5 | **$7.95\text{x}10^{-11}$** |
| Biological entity * History * Environment | 4 | 47.9 | **$1.01\text{x}10^{-9}$** |
| | | *$R^2$=0.85* | |
